# High performance *Legionella pneumophila* source attribution using genomics-based machine learning classification

**DOI:** 10.1101/2023.03.19.532693

**Authors:** Andrew H. Buultjens, Koen Vandelannoote, Karolina Mercoulia, Susan Ballard, Clare Sloggett, Benjamin P. Howden, Torsten Seemann, Timothy P. Stinear

**Affiliations:** Department of Microbiology and Immunology, Doherty Institute for Infection and Immunity, University of Melbourne, Melbourne, Victoria, Australia; Centre for Pathogen Genomics, University of Melbourne, Melbourne, Victoria, Australia; Bacterial Phylogenomics Group, Institut Pasteur du Cambodge, Phnom Penh, Cambodia; Microbiology Diagnostic Unit, Department of Microbiology and Immunology, Doherty Institute for Infection and Immunity, University of Melbourne, Melbourne, Victoria, Australia; Department of Infectious Diseases, Austin Health, Heidelberg, Victoria, Australia

## Abstract

Fundamental to effective Legionnaires’ disease outbreak control is the ability to rapidly identify the environmental source(s) of the causative agent, *Legionella pneumophila*. Genomics has revolutionised pathogen surveillance but *L. pneumophila* has a complex ecology and population structure that can limit source inference based on standard core genome phylogenetics. Here we present a powerful machine learning approach that assigns the geographical source of Legionnaires’ disease outbreaks more accurately than current core genome comparisons. Models were developed upon 534 *L. pneumophila* genome sequences, including 149 genomes linked to 20 previously reported Legionnaires’ disease outbreaks through detailed case investigations. Our classification models were developed in a cross-validation framework using only environmental *L. pneumophila* genomes. Assignments of clinical isolate geographic origins demonstrated high predictive sensitivity and specificity of the models, with no false positives or false negatives for 13 out of 20 outbreak groups, despite the presence of within-outbreak polyclonal population structure. Analysis of the same 534-genome panel with a conventional phylogenomic tree and a core genome multi-locus sequence type allelic distance-based classification approach revealed that our machine learning method had the highest overall classification performance – agreement with epidemiological information. Our multivariate statistical learning approach maximises use of genomic variation data and is thus well-suited for supporting Legionnaires’ disease outbreak investigations.

## INTRODUCTION

*Legionella pneumophila* is a gram negative bacterium that can thrive in warm, moist built environments and then cause Legionnaires’ disease (LD) in humans when contaminated water is aerosolised and inhaled (David et al., 2016; Fields, Benson, & Besser, 2002; Mercante & Winchell, 2015; Schwake, Garner, Strom, Pruden, & Edwards, 2016). The vast majority of clinical infections are caused by *L. pneumophila* serogroup 1 (Yu et al., 2002). To combat LD outbreaks, public health authorities must rapidly investigate and determine the environmental sources to then intervene to prevent further disease transmission. A major difficulty in pin-pointing source(s) is the fact that there often exist a multitude of possible origins, particularly in densely populated urban settings.

The advent of bacterial genotyping has been advantageous for LD outbreak investigations, helping to ‘rule in’ or ‘rule out’ suspected environmental sources by attempting to match the genotypes of *L. pneumophila* recovered from patients to those derived from a given environmental source. In particular, Sequence Based Typing (SBT) compares DNA sequence variations across seven core genes to generate a sequence type (ST) that is standardised and internationally recognised (Lück, Fry, Helbig, Jarraud, & Harrison, 2013). An ST can be used to assign isolates from clinical specimens to specific environmental sources. Despite its popularity and simple interpretation, the SBT scheme lacks discriminatory power. The scheme captures only a tiny fraction of bacterial genomic variation and this is problematic when the majority of LD cases are caused by just a handful of STs (Borchardt, Helbig, & Lück, 2008; David et al., 2016; Harrison, Afshar, Doshi, Fry, & Lee, 2009). SBT is thus largely inadequate for LD source investigations.

Whole genome sequencing is used increasingly routinely for public health surveillance and infectious disease outbreak investigations and recent efforts have utilised the power of genomics to confirm suspected bacterial pathogen environmental sources (Abrams & Trees, 2017; Goldberg, Sichtig, Geyer, Ledeboer, & Weinstock, 2015; Krøvel et al., 2022; Petzold, Prior, Moran-Gilad, Harmsen, & Lück, 2017; Ricci et al., 2022; Rousseau et al., 2022; Schoonmaker-Bopp et al., 2021; Wüthrich et al., 2019). In particular, genomic analyses that assess core-genome variation (sites present in all isolate genomes) such as phylogenomic trees and pairwise SNP distances, have been useful to investigate disease transmission (Gorrie et al., 2021; Ingle, Howden, & Duchene, 2021; Kwong et al., 2016; Sintchenko & Holmes, 2015).

Another genomics-based approach for *L. pneumophila* source tracking is the core genome multi locus sequence typing (cgMLST) scheme that builds upon the SBT concept but greatly expands the genomic variation that is considered (Moran-Gilad et al., 2015). In cgMLST, the allele scheme is enlarged from seven core genes to a panel of 1,521 genes to produces an allele-type integer for each novel variant combination (Moran-Gilad et al., 2015). This systematised and expanded approach provides greater discrimination compared to conventional SBT, however like phylogenomic approaches, it is still limited to only core-genome variation. Despite the increased utility of such core-genome based approaches compared with SBT, they still lack adequate discriminatory power for investigation of some *L. pneumophila* outbreaks where isolate genomes are often near identical at the core-genome level (Buultjens et al., 2017; McAdam et al., 2014; Sánchez-Busó et al., 2016).

An alternative to core-genome analyses is to incorporate variation in accessory genome sites; that is, to use DNA sequences present in some but not all isolates. Here, to make better use of all the available genomic variation, we have developed a machine learning statistical modelling method that utilises SNP variation in both the accessory and core genome (pan-genome SNP variation) to classify genomes by likely environmental source. Our approach integrates pan-genome SNP variation using multivariate algorithms that model interrelationships among multiple variables to assign source with greater accuracy than a standard core-genome SNP comparison approach. This advance builds on our previously reported *L. pneumophila* source tracking modelling approach that had high positive classification capacity (rule-in) but had no negative classification ability (rule-out) (Buultjens et al., 2017).

In this study, we have implemented ‘one-versus-rest’ machine learning classifier algorithms with the ability to reject *L. pneumophila* clinical isolate genomes that don’t belong to classes used to train models, achieving both high classification sensitivity and specificity. We have benchmarked the classification performance of our machine learning method against phylogenomic and cgMLST allele distance approaches using epidemiological assignments. Our machine learning algorithms built with pan-genome SNP variants allowed us to assign the environmental sources of LD outbreaks and make objective assignments of clinical isolate genome origins. It is envisioned that future LD public health investigations may make use of such sensitive and specific multivariate modelling advancements to rapidly identify the environmental source of *L. pneumophila* and reduce the spread of this preventable disease.

## METHODS

### Bacterial genomes used in this study

The isolate genomes originating from this study were cultured and sequenced as per previously described (Buultjens et al., 2017). WGS data for an international collection of diverse *L. pneumophila* (spanning 23 STs) was included in this study (Supplementary Table. S1). A total of 246 isolates in this study were newly sequenced while 288 were publicly available as either draft genome assemblies or raw reads.

### Reference based core genome SNP calling

Snippy v4.4.5 was used to map reads and contigs to a previously described fully assembled *L. pneumophila* clinical isolate genome Lpm7613 originating from Melbourne, Australia (GenBank assembly accession: GCA_900092465.1) using a ‘minfrac’ setting of 0.8 (https://github.com/tseemann/snippy). The snippy-core subcommand was used to generate a core genome SNP alignment - SNP variation in the fraction of the genome shared by all isolates. Pairwise SNP differences were assessed using a custom R script (https://github.com/MDU-PHL/pairwise_snp_differences).

### Phylogenomic tree analysis

Clonal Frame ML was used to infer sites impacted by recombination (Didelot & Wilson, 2015). The regions predicted to have been affected by recombination were used to generate a bed file that was subsequently used for masking of the core genome alignment with Snippy (see above). A maximum likelihood phylogenomic tree was built from the alignment of non-recombining core SNPs using FastTree v2.1.10 (Price, Dehal, & Arkin, 2009). Trees were displayed using FigTree v1.4.4 (http://tree.bio.ed.ac.uk/software/figtree). The cophenetic function of the ape R package v5.6-2 (Paradis, Claude, & Strimmer, 2004) was used to compute a 534 × 534 patristic distance matrix from the tree newick file.

### Core genome Multi Locus Sequence Typing

Core genome MLST analysis was undertaken using Coreugate v2.0.5 (https://github.com/kristyhoran/Coreugate). Draft genome assemblies were generated by shovill v0.9.0 (https://github.com/tseemann/shovill) using the SPAdes genome assembler v3.15.2 (Bankevich et al., 2012) and provided as input for Coreugate (filter_samples_threshold=0.85). The *L. pneumophila* allele scheme used with Coreugate was described by (Moran-Gilad et al., 2015).

### Reference independent pan genome SNP calling

Split Kmer Analysis (SKA) v1.0 was used to detect pan-genome SNPs (SNPs in core and accessory sites) from reads and assembly contigs (Harris, 2018). Raw reads were trimmed of adapter sequences using Trimmomatic v0.39 using the ‘-phred33’ option (Bolger, Lohse, & Usadel, 2014). Here, the fastq and fasta subcommands were used to generate split kmer files (kmer size of 15) from isolates with reads in the fastq file format and assembly contigs in fasta format, respectively. The split kmer files were combined using the align subcommand (p=0.1) to produce a reference-independent pan-genome SNP alignment and the humanise subcommand was used to generate a SNP matrix from the skf alignment file.

### Distance-based classification

Matrices of pairwise distances were generated from both the phylogenomic tree and the cgMLST alleles and used to devise distance-based classifiers. The average distance among the environmental isolates for each outbreak group was calculated and used as the outbreak group specific cut-off threshold to then classify the 113 clinical isolate genomes as either being outbreak related or not. This analysis was conducted only for outbreak groups that had at least two environmental isolate genomes available (14 of the 20 groups) (Table. 1).

**Table 1.** Attributes of the 534 L. pneumophila isolates included in this study.

| Group | Number of<br>environmental<br>isolates | Number of<br>clinical<br>isolates | STs | Reference |
| --- | --- | --- | --- | --- |
| ENOA | 311 | NA | 15 SBTs | This study; Bartley, PB., et. al., 2016; Buultjens, AH., et. al., 2017; David, S., Rusniok, C., et. al., 2016; David, S., et. al., 2017; Moran-Gilad, J., et. al., 2015; Qin, T., et.al., 2016 |
| CNOA | NA | 74 | 9 SBTs | Bartley, PB., et. al., 2016 ; Buultjens, AH., et. al., 2017 ; David, S., Rusniok, C., et. al., 2016; David, S., et. al., 2017; Moran-Gilad, J., et. al., 2013 |
| MELB-2018 | 20 | 3 | SBT30 | This study |
| MELB-A | 14 | 11 | SBT30 | Buultjens, AH., et. al., 2017 |
| MELB-C | 3 | 1 | SBT30 | Buultjens, AH., et. al., 2017 |
| MELB-G | 18 | 2 | SBT30 | Buultjens, AH., et. al., 2017 |
| MELB-M | 8 | 1 | SBT30 | Buultjens, AH., et. al., 2017 |
| ESSEX-A | 7 | 2 | SBT1 | David et al., 2017 |
| ESSEX-B | 3 | 1 | SBT1 | David et al., 2017 |
| ESSEX-E | 2 | 1 | SBT1 | David et al., 2017 |
| ESSEX-G | 14 | 1 | SBT1 | David et al., 2017 |
| ESSEX-H | 5 | 2 | SBT1 | David et al., 2017 |
| NY-1 | 3 | 1 | SBT1 | Raphael, B. Baker, D., et. al., 2016 |
| NY-2 | 2 | 2 | ND | Raphael, B. Baker, D., et. al., 2016 |
| NY-3 | 1 | 3 | SBT1 | Raphael, B. Baker, D., et. al., 2016 |
| NY-4 | 1 | 1 | SBT1 | Raphael, B. Baker, D., et. al., 2016 |
| NY-5 | 1 | 1 | SBT62 | Raphael, B. Baker, D., et. al., 2016 |
| NY-6 | 3 | 1 | SBT36 | Raphael, B. Baker, D., et. al., 2016 |
| NY-7 | 1 | 1 | SBT36 | Raphael, B. Baker, D., et. al., 2016 |
| NY-8 | 1 | 1 | SBT1204 | Raphael, B. Baker, D., et. al., 2016 |
| NY-9 | 2 | 1 | SBT94 | Raphael, B. Baker, D., et. al., 2016 |
| NY-10 | 1 | 2 | SBT731 | Raphael, B. Baker, D., et. al., 2016 |
| **Total** | **421** | **113** |  |  |

### Classifier evaluation

Performance of all classifiers was assessed using the F1 metric. The F1-score is the harmonic mean of the recall and precision, conveying the balance between these two metrics. Here, a F1-score of 1 indicates that the classifier performs perfectly (no false positives or false negatives). The F1-score is particularly useful to appraise classification models when there is class imbalance.

### Machine learning classification framework

#### Preparation of test and train datasets

The SKA pan-genome SNP matrix was one-hot encoded using the scikit-learn library pre-processing module (Géron, 2019). The encoded matrix was divided into separate training and testing datasets upon whether the isolate genomes were sourced from either environmental samples, for training (n=421), or clinical samples, for testing (n=113) (Fig. 1A).

**Fig. 1.**
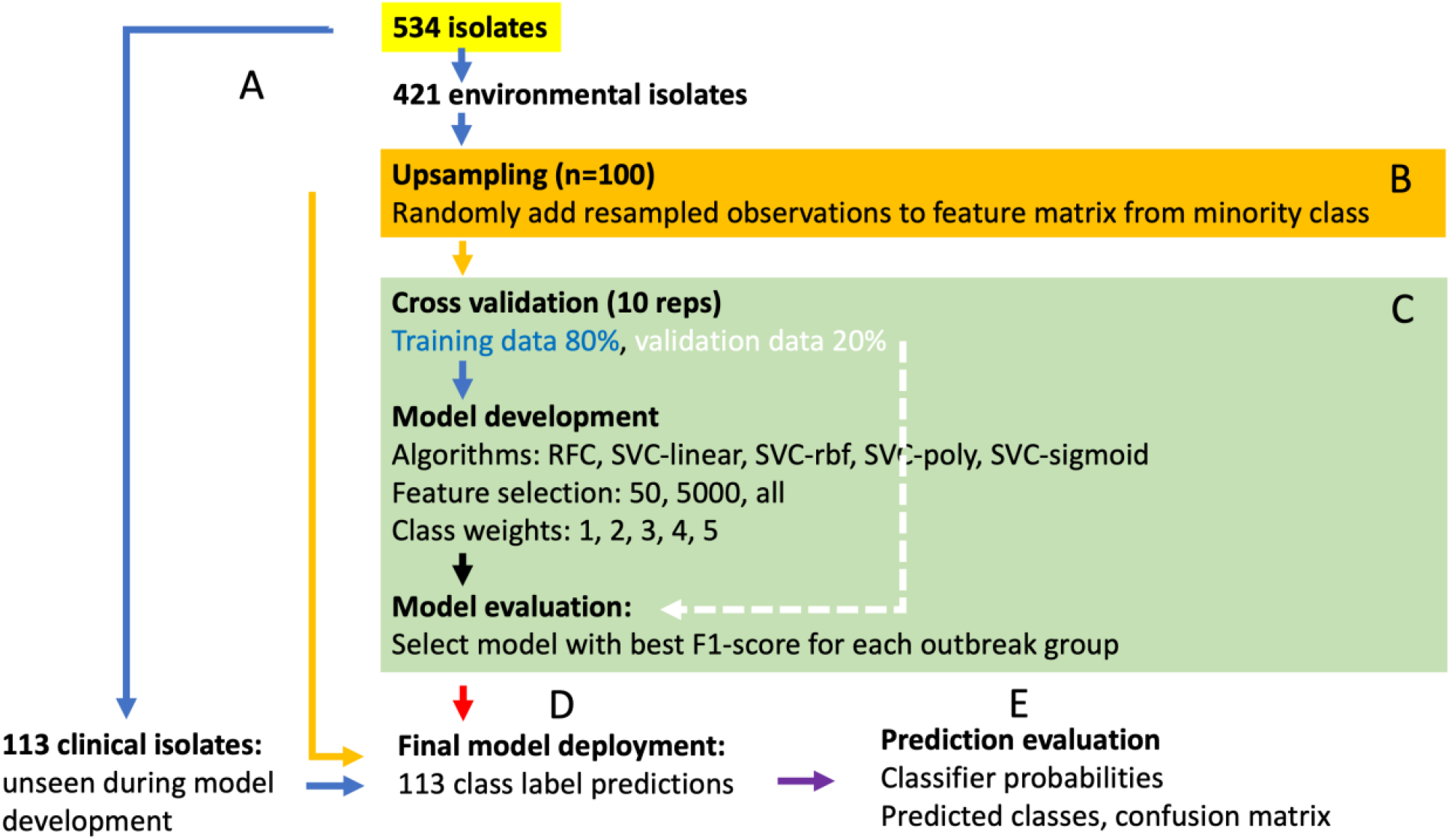
Flow diagram of the machine learning model development framework. A) Isolate genomes were separated from the input one-hot encoded matrix (n=534) according to being either environmentally (n=421) or clinically derived (n=113). B) Upsampling was performed on the environmental training dataset, where individuals from the minority class were randomly upsampled (with replacement). C) Cross validation loop. In each iteration, the training data was randomly split into a training and testing partition of 80% and 20%, respectively, for ten repetitions. Various combinations of model parameters were used, and the classifier was evaluated upon ability to correctly assign the test set component of the data using the F1-score. D) The models for each outbreak group with the greatest F1-score in the cross- validation loop were selected to form a set of final models. Final models were trained with all available upsampled environmental isolate genome data to then assign the classes of the previously unseen 113 clinical isolate genomes. E) Classification outputs were in the form of probabilities that were binarised as either belonging to or not belonging to each specific outbreak group class. Information of clinical isolate known origins was used to establish a confusion matrix and calculate the F1-score.

#### Model development

As the available epidemiological information was discrete geographical locations, a supervised classification approach was used. Here the class labels were formatted to represent a binary array of ‘1’ (linked to outbreak) and ‘0’ (not linked to outbreak). The use of separate label files for each outbreak cluster allowed for the implementation of a ‘one-vs-rest’ classification framework, in which each outbreak group had its own model built, with the learning objective to include isolates of class ‘1’ and reject those of class ‘0’.

### Upsampling to redress class imbalance

Due to the availability of few outbreak-associated environmental isolate genomes compared to that of clinical isolate genomes, there existed a substantial imbalance between the ‘0’ and ‘1’ classes in the training set. To reduce this class imbalance, *upsampling* was implemented in which observations from the minority class were randomly selected (with replacement) and appended to the feature matrix (Fig. 1B). As each outbreak group had a different set of labels, this was undertaken for all 20 outbreak groups. Given that there were approximately 20 minority class environmental isolate genomes to 400 majority class observations, an upsampling amount of 100 was chosen as this was approximately 1/4 of the majority class in each situation - a conservative upsampling portion given the severe class imbalance. The remaining class imbalance was addressed through specifying class weights to the classification algorithm (see below).

### Model development cross-validation

During model training, supervised classification algorithms learn specific patterns associated with each of the classes with the goal to develop models that are generalisable, in that they can make accurate assignments upon previously unseen observations. To promote optimal model development on the environmental isolate training dataset, an iterative cross-validation procedure was undertaken to determine the best model for each outbreak group (Fig. 1C). Here, the training data was randomly split into training and validation partitions (80% train and 20% validation) 10 times, with models built upon the training portion and used to classify the classes of the validation portion. For each iteration in the cross-validation loop, the F1-score was recorded and used to evaluate each model. A different set of model parameter combinations was evaluated with each cross-validation iteration (model parameters: classifier algorithm, class weights, and number of selected features) (Fig. 1C). A total of 1,500 model combinations were evaluated in the cross-validation phase.

### Multivariate classification algorithms

Two supervised classifier algorithms were implemented: Random Forest Classifiers (RFC) and Support Vector Classifiers (SVC) (Fig. 1C). RFC indiscriminately select a subset from the training data to create a collection of decision tree predictors to sum the predictions, in effect lowering the variance (Breiman, 1996). Here, each decision tree takes a set of features and provides an individual output, all of which are subsequently summarised to produce a final probabilistic output (Breiman, 2001). The scikit-learn RFC module was implemented with default parameters (Géron, 2019). SVC optimise for non-linear combinations of features that best divide the classes across a multi-dimensional hyperplane (Boser, Guyon, & Vapnik, 1992). The scikit-learn SVC module was implemented with default parameters apart from using kernels: ‘linear’, ‘rbf’, ‘poly’ and ‘sigmoid’ (Géron, 2019).

### Class weights

A further approach to combat the occurrence of class imbalance was to specify class weights to the classification algorithms. The reasoning here was that classifiers have default assumptions of class balance and, when faced with class imbalance, a bias exists that favours towards the dominant class. In this case the ‘0’ or ‘not outbreak related’ isolates are likely to cause bias, as they strongly outnumber the amount of ‘1’ or ‘outbreak related’ isolates. By specifying class weights the classification algorithm is modified to account for the skewed class distribution, enabling improved training and higher performance assignments by penalising misclassification of the minority class. Specifically, the class weights were passed to the scikit-learn classifiers as a dictionary that stipulated class ‘0’ as 0.5 and class ‘1’ as an integer in the range of 1 to 5 (Fig. 1C).

### Univariate feature selection

Features that did not vary in proportion between the classes for a particular outbreak group are unlikely to have any classification value for model training and therefore only add noise. To reduce the number of uninformative features and focus on those that are associated with the class labels, feature selection was performed. The SelectKBest univariate module of scikit-learn was employed to assess the independence of individual features against the target variable using a chi-square test, selecting the top 50, 5,000 or all features (Fig. 1C) (Géron, 2019). To avoid any data-leakage, the univariate feature selection was only performed on the training set either during the cross-validation procedure or on all available environmental isolates for the building of the final models (see below).

### Final model classifications

Following selection of the top performing model combinations for each outbreak group, final models were built using the model parameters identified and trained with all available environmental isolates (n=421). Here, the final model for each outbreak group learned as much as possible about the genomic variability in the data when all available environmental isolates were used (Fig. 1D-E). Thus, this was the optimal way to train the final models to make generalisable source attribution assignments upon the clinical isolate genomes.

The code used to conduct the abovementioned analyses is detailed in the following github repository: https://github.com/abuultjens/Assign_Legionella_pneumophila_origins

## RESULTS

### Selection of *L. pneumophila* genome sequences for classification model development

The overall objective of this research was to attempt to use multivariate statistical learning methods to assign the environmental sources of LD outbreaks. However, to benchmark the performance of such methods it was first necessary to select a set of *L. pneumophila* genomes representing different LD outbreak investigations. Our principles for genome selection were to maximise both genomic and spatial diversity to achieve a collection that spanned many Sequence Types (STs) and originated from various locations worldwide. A review of the literature and publicly available *L. pneumophila* genome sequences revealed studies from three different jurisdictions (see details below) spanning 20 distinct LD outbreaks that were suitable to include because they had sufficiently rich epidemiological information and associated *L. pneumophila* genomic data from both clinical and environmental sources. In all, 534 *L. pneumophila* genomes were identified for use in this study, of which 421 and 113 represented bacterial isolates from environmental and clinical sources, respectively (Table. S1).

The outbreak associated group consisted of 149 isolates that were epidemiologically linked to a total of 20 outbreaks across three major geographical regions: 1) Melbourne, Victoria, Australia, 2) Essex, England, and 3) New York State, United States (Table. S1). The Melbourne *L. pneumophila* genomes, hereon referred to with prefix “MELB”, represented five different LD outbreaks spread across the Melbourne metropolitan area, occurring between 1998-2018 (Buultjens et al., 2017). The Essex *L. pneumophila* genomes, hereon referred to with prefix “ESSEX”, consisted of genomes obtained from *L. pneumophila* isolates linked to LD disease occurring in five distinct wards within a single hospital campus (isolated between 2007-2011) (David et al., 2017). The New York State *L. pneumophila* genomes, hereon referred to with prefix “NY”, consisted of 10 separate LD outbreaks across the New York State area (*L. pneumophila* isolated between 2004-2012) (Raphael et al., 2016).

To assist in developing a classification framework with negative classification capacity, *i.e.* the ability of the model to call true negatives, we included genome sequences from 74 *L. pneumophila* clinical isolates not associated with any of the abovementioned outbreaks, hereon referred to as clinical non-outbreak associated (CNOA) (Table. 1). These isolate genomes were isolated between 1986-2014 and originated from across Europe, the United Kingdom and Australia. In a similar way, to challenge the model building process, we included 311 environmental isolates (isolated between 1995-2018) that were not associated with any of the outbreaks (MELB, ESSEX or NY), hereon referred to as the environmental non-outbreak associated (ENOA) (Table. 1).

### Population structure of *L. pneumophila* isolates used in this study

We examined the genomic context of 421 environmental *L. pneumophila* isolate genomes alongside 113 clinical isolate genomes to investigate the ability to make inferences of source attribution. The 20 outbreak groups were from three distinct geographical regions, Melbourne (Australia), Essex (UK) and New York (US). Sequence read alignment against a SBT30 reference genome revealed 221,214 core genome SNPs. There were 144,829 SNP sites inferred to have arisen by recombination, leaving 76,385 SNPs that were derived through vertical transmission. Pairwise SNP comparisons were performed to depict the amount of diversity within each outbreak group (Fig. 2A). Most of the groups had mean intra-group distances between 0-4 SNPs, while MELB-A, NY-3 and NY-8 had elevated within group variations of 41, 61 and 24 SNPs, respectively.

**Fig 2.**
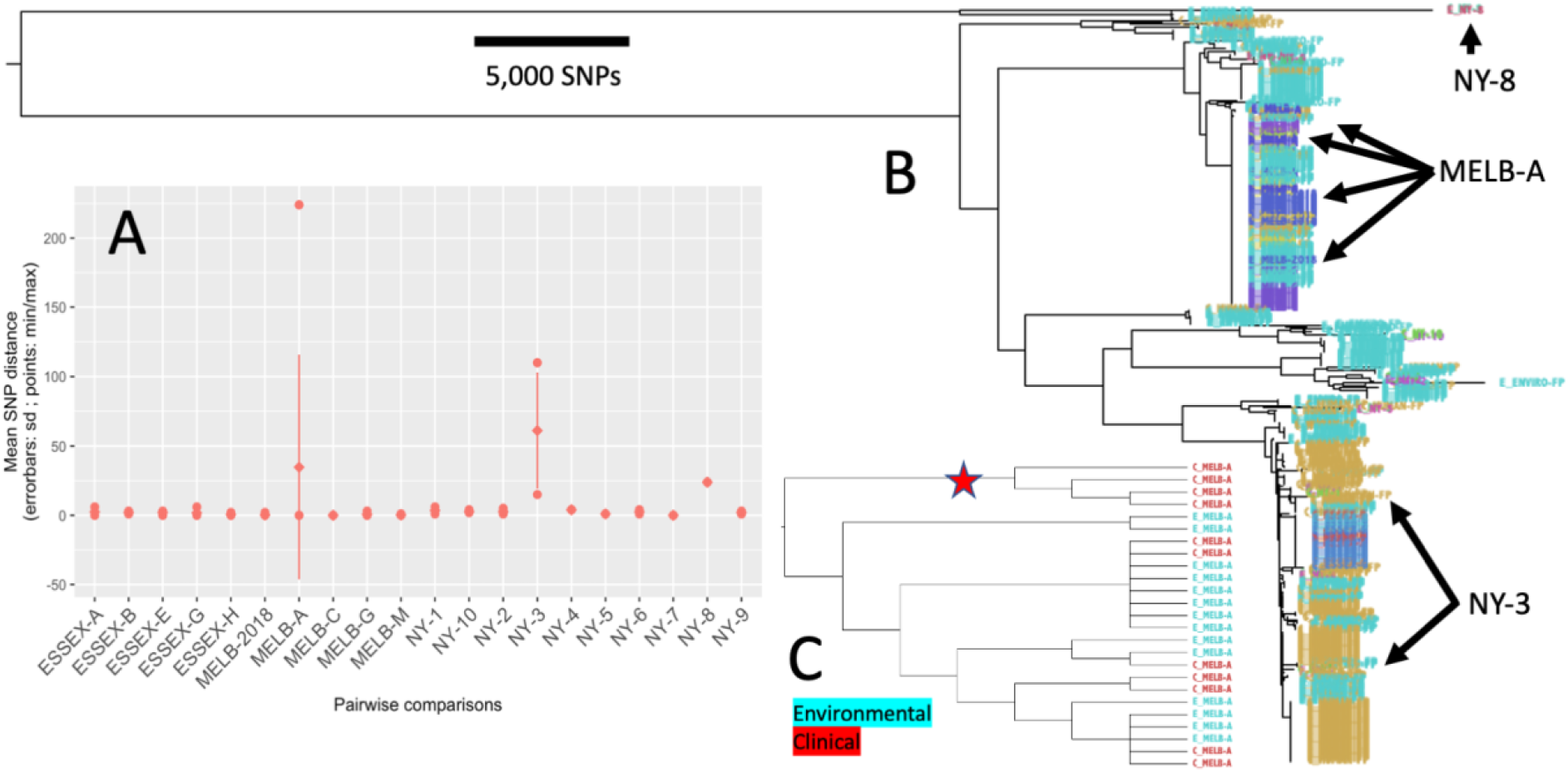
Assessment of genomic population structure of 534 *L. pneumophila* clinical and environmental isolate genomes. A) Pairwise SNP comparisons of within outbreak group diversity. Three groups had elevated levels of within group diversity: MELB-A, NY-3 and NY-8. B) Phylogenomic tree generated from non-recombining core genome SNPs. Outbreak groups MELB-A, NY-3 and NY-8 are indicated C) Subtree containing isolate genomes associated with the MELB-A outbreak. The subtree is displayed as a cladogram with branch lengths transformed to illustrate the tree topology. Red star indicates a distinct genotype containing only clinical isolate genomes without any environmental representatives.

Phylogenomic analysis has become an important approach to examine pathogen population structure and to investigate the likely origins of *L. pneumophila* clinical isolates using WGS data (David et al., 2016; Gorzynski et al., 2022; Graham, Doyle, & Jennison, 2014; Qin et al., 2016; Reuter et al., 2013; Wüthrich et al., 2019). A phylogenomic tree was estimated from the non-recombining core-genome SNP alignment to depict the clonal ancestry (Fig. 2B). The tree illustrated the same grouping of outbreak related isolates that was observed with the pairwise SNP distance analysis. In particular, the groups with high internal SNP diversity displayed the existence of within-outbreak polyclonal population structure (Fig. 2B). The MELB-A isolate genomes were found to harbor several distinct genotypes, one of which was exclusively represented by clinical isolates (Fig. 1B-C). Outbreak group NY- 3 isolate genomes were located across several distinctive subtrees in the phylogeny, indicating a within group polyclonal population structure (Fig. 1B). NY-8 isolate genomes had an elevated within group diversity while also being substantially distinct to all other isolates included in the study (Fig. 1B).

### Phylogenomic tree distance-based classification

To objectively assess the ability to infer clinical isolate origins from the phylogenomic tree, patristic distances were extracted and used to build outbreak group specific classifiers. Here, the patristic distances represent the individual total branch length distances between all possible isolate pairs in the tree, represented as a 534 x 534 distance matrix. The average distance between the environmental isolate genomes of each outbreak group were calculated for groups that had at least two or more environmental representatives (14 of the 20 outbreak groups). The average distance among environmental isolate genomes was used as a threshold to assign each query clinical isolate as either related or unrelated to the outbreak groups, with each group having a specific threshold distance (14 different thresholds and classifiers). The assumption underlying the use of distance thresholds was that a clinical isolate genome with equal or less patristic distance from the mean distance observed among environmental isolate genomes from a specific outbreak group is likely related to that outbreak while those with greater distances are more divergent and thus likely originated elsewhere.

The cut-off distance threshold for each outbreak group was determined through analysis of only the environmental isolate genomes for each specific outbreak group. This is an ideal approach, as the thresholds are not biased by the addition of any clinical isolate genomes, therefore building a classification tool that is prospective, in that the system would be ready for deployment before the first clinical isolate genome is reported in an outbreak investigation. The performance of the classifiers was assessed using the F1-score which is the harmonic mean of method recall and precision, conveying the balance between these two metrics. Here, a F1-score of 1 indicates that the classifier performs perfectly (no false positives or false negatives). The patristic distance-based classifiers demonstrated the ability to correctly assign most clinical isolate genomes to their known origins (0.43 mean false negatives), however this approach had a high false positive rate (3.93 mean false positives) with an overall mean F1 score of 0.50 (Table. 2).

**Table 2.** Distance-based and machine learning classifications for 113 test set clinical isolate genomes when trained on a set of 421 training environmental isolate genomes.

|  | Patristic distance-based classifiers |  |  | cgMLST distance-based classifiers |  |  | Machine learning classifiers |  |  |
| --- | --- | --- | --- | --- | --- | --- | --- | --- | --- |
| Outbreak group | False positive | False negative | F1-score | False positive | False negative | F1-score | False positive | False negative | F1-score |
| MELB-2018 | 4 | 0 | 0.60 | 2 | 0 | 0.75 | 0 | 0 | 1 |
| MELB-A | 6 | 4 | 0.58 | 18 | 0 | 0.55 | 1 | 5 | 0.67 |
| MELB-C | 11 | 0 | 0.15 | 6 | 0 | 0.25 | 0 | 1 | 0 |
| MELB-G | 7 | 0 | 0.36 | 19 | 0 | 0.17 | 0 | 0 | 1 |
| MELB-M | 11 | 0 | 0.15 | 22 | 0 | 0.08 | 0 | 0 | 1 |
| ESSEX-A | 5 | 0 | 0.44 | 2 | 1 | 0.40 | 1 | 1 | 0.5 |
| ESSEX-B | 4 | 0 | 0.33 | 3 | 0 | 0.4 | 1 | 1 | 0 |
| ESSEX-E | 4 | 0 | 0.33 | 6 | 0 | 0.25 | 0 | 1 | 0 |
| ESSEX-G | 3 | 0 | 0.40 | 4 | 0 | 0.33 | 1 | 0 | 0.67 |
| ESSEX-H | 0 | 1 | 0.67 | 0 | 1 | 0.67 | 0 | 0 | 1 |
| NY-1 | 0 | 0 | 1 | 0 | 0 | 1 | 0 | 0 | 1 |
| NY-2 | 0 | 0 | 1 | 0 | 0 | 1 | 0 | 0 | 1 |
| NY-3 | NA | NA | NA | NA | NA | NA | 0 | 3 | 0 |
| NY-4 | NA | NA | NA | NA | NA | NA | 0 | 0 | 1 |
| NY-5 | NA | NA | NA | NA | NA | NA | 0 | 0 | 1 |
| NY-6 | 0 | 0 | 1 | 0 | 0 | 1 | 0 | 0 | 1 |
| NY-7 | NA | NA | NA | NA | NA | NA | 0 | 0 | 1 |
| NY-8 | NA | NA | NA | NA | NA | NA | 0 | 0 | 1 |
| NY-9 | 0 | 1 | 0 | 0 | 0 | 1 | 0 | 0 | 1 |
| NY-10 | NA | NA | NA | NA | NA | NA | 0 | 0 | 1 |
| AVERAGE | *3.93 | *0.43 | *0.50 | *5.86 | *0.14 | *0.56 | 0.20<br>(*0.29) | 0.60<br>(*0.64) | 0.74<br>(*0.70) |

\* When considering groups with two or more environmental isolate genomes

### cgMLST distance-based classification

In addition to phylogenomics, cgMLST is another genomic comparison approach used to infer the source attribution of *L. pneumophila* clinical isolate genomes which builds upon the established SBT genotyping method by greatly expanding the number of core-genome loci (Moran-Gilad et al., 2015; Qin et al., 2016). The advantage of cgMLST over analyses that consider all core genome SNPs is the standardised framework in which the alleles are called, in that cgMLST is not susceptible to fluctuations in core genome size caused by the addition or removal of isolates from the analysis. We next investigated if the allelic distance derived from the cgMLST scheme, when applied to the 534 isolates, could be used to provide improved source attribution inference. Here, the same threshold derivation and classification approach that was employed for the patristic distances was applied, however using a distance matrix generated from cgMLST allelic variation.

The cgMLST based classifiers had fewer false negatives than the patristic distance-based classifiers (0.14 mean false negatives) while having a higher false positive rate (5.86 mean false positives) and a marginally higher overall mean F1-score of 0.56 (Table. 2). The classifiers performed well for NY outbreak groups that had more than one environmental isolate genome, all achieving F1-scores of 1. While the implementation of phylogenomic tree and cgMLST distance-based classifiers introduced an objective framework to make source inferences, these approaches were based solely on core-genome variation, raising the question of whether approaches built using SNP variation from across the pan-genome may achieve greater assignment capacity.

### Machine learning classification

To enhance the classification capacity of the framework, we applied a machine learning approach that utilised an alignment containing 479,480 SNPs detected in both core and non-core sites. The advantage of using pan-genome SNPs for this type of analysis was that additional variation in accessory genome sites is thus considered, improving the discriminatory potential for downstream analyses. In addition to greater SNP variation, the use of a multivariate classification algorithm provides the advantage in that the concerted effects of all input genomic variants are modelled to learn about informative structures in the data.

To reduce the likelihood of overfitting, a cross-validation framework was established that iteratively split the environmental isolate data into train and validation partitions. A total of 1,500 model combinations consisting of different model parameters using both Random Forest Classifiers (RFC) and Support Vector Classifiers (SVC) (see methods) were evaluated. In this way, the best model combination for each outbreak group was determined using only environmental isolate genomic variation prior to the analysis ever encountering any clinical isolate genomes, thus eliminating the risk of model overfitting, and providing a prospective approach.

### Machine learning model results

Application of the final models for the assignment of the clinical isolate genomes provided the lowest false positive rate of all previous distance-based approaches (0.29 mean false positives), the highest level of false negatives (0.64 mean false negatives) and the highest overall mean F1 score of 0.70 when applied to the 14 outbreaks with two or more environmental isolate genomes (Table. 2). Models developed for outbreak groups MELB-2018, MELB-G, MELB-M, ESSEX-H, NY-1 through NY-2 and NY-4 through NY-10 (13/20) had F1-scores of 1, indicating the absence of any false positives or false negatives – classifications that perfectly align with the epidemiological labels (Table. 2). As the machine learning method used upsampling to artificially replicate the training observations, it was possible to apply this method to outbreak groups with as few as one environmental isolate genome, having an overall mean F1-score of 0.74 when applied to all 20 outbreak groups. (Table. 2). The parameters of the final models are reported in Supplementary Table 2.

### Examination of machine learning model false positives and false negatives

False positives occurred with models ESSEX-A, ESSEX-B and ESSEX-G. In these instances, the false positives were from other ESSEX outbreak groups clinical isolate genomes (wards within the same hospital). Six of the models MELB-A, MELB-C, ESSEX-A, ESSEX-B, ESSEX-E and NY-3 had one or more false negative classifications. In the case of NY-3, there was an appreciable amount of within outbreak diversity (Fig. 2A) and just a single environmental isolate used for model training (Table. 1). For MELB-A, the four clinical isolates that were classified as false negatives by the machine learning approach were on a branch in the phylogeny that did not contain any MELB-A environmental isolate genomes and therefore were from a specific genotype that was not represented in the training data (Fig. 2C). Despite this, all 11 of the MELB-A clinical isolate genomes were within the top 21% of the 113 test-set clinical isolate genomes when ranked according to decreasing classification probability (Fig. 3A).

**Fig 3.**
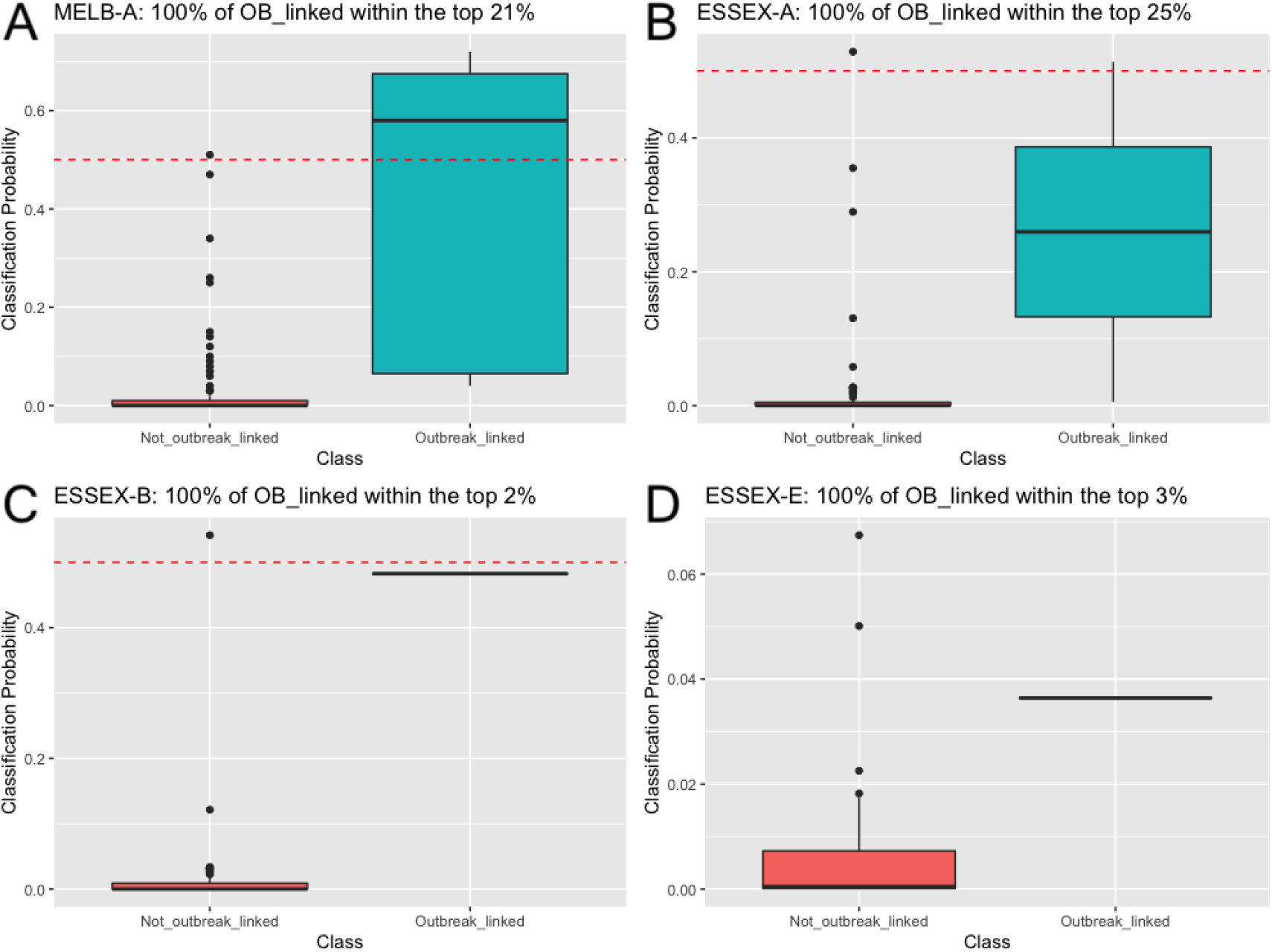
Boxplots of classification probabilities for the outbreak linked and non-outbreak linked 113 test set clinical isolate genomes for four outbreak group models that had false negative classifications. Red horizontal dotted lines indicate the classification threshold of 0.5. A: classification probabilities for outbreak group MELB-A. B: classification probabilities for outbreak group ESSEX-A. C: classification probabilities for outbreak group ESSEX-B. D: classification probabilities for outbreak group ESSEX-E.

Investigation of the machine learning classifier probabilities for outbreak groups ESSEX-A, ESSEX-B and ESSEX-E also revealed that despite having false negatives at the default classification threshold of 0.5, the classification probabilities were nonetheless informative to rank the clinical isolate genomes (Fig. 3B-D). In this way, when ranked according to decreasing probabilities, the clinical isolate genomes from ESSEX-A, ESSEX-B and ESSEX-E were contained withing the top 25%, 2% and 3% of all clinical isolate genomes, respectively (Fig. 3B-D). In these instances, if the classification threshold were lower than 0.5, these models would have provided perfect and near perfect classifications.

## DISCUSSION

Timely and accurate identification of environmental sources of LD is of utmost importance to public health investigations and, in this era of high-resolution genomic technologies, innovative approaches are needed to rapidly distil complex analyses to provide actionable insights. In this study, we have deployed a machine learning classification approach and assessed its’ ability alongside alternative approaches to make assignments of clinical isolate origins that align against the known epidemiological information for 20 distinct LD outbreaks.

This work builds on our previous efforts to build accurate multivariate assignment models, here providing the necessary negative classification capacity that was lacking in our earlier work. To assess the ability of these multivariate approaches to call true negatives, we included 74 clinical isolates that were not associated with any of the 20 outbreak groups that were used to train the models. In a similar way, we also included 311 environmental isolates that were not associated with the outbreak groups to assess how well the model could learn from known outbreaks while faced with a larger than necessary training dataset that contained unrelated environmental isolates. Our improved approach presented in this investigation made use of a set of ‘one-vs-rest’ classification strategies, in which a separate target variable and model was used for each outbreak group. This had the effect of focusing on genomic variation that was specific to an individual outbreak group, optimising the model to include outbreak linked isolates while rejecting others and therefore affording negative classification capacity.

The analysis of suspected pathogen transmission with phylogenomic trees built from core genome SNPs has become the *de facto* standard in the field of bacterial genomics. Here we assessed the ability of patristic distances derived from a phylogenomic tree to place epidemiologically linked isolate genomes into arrangements that could then permit the inference of clinical isolate source attribution. Classifiers were devised for the 14 of the 20 groups that had at least two environmental isolate genomes, with assignment thresholds derived from the mean distance observed among the environmental representatives of each group. This approach provided an objective and quantitative phylogenomic-based framework for the classification of query clinical isolate genomes with high sensitivity; however, it suffered from low specificity and had an overall mean F1-score of 0.50.

Another widely employed tool for *L. pneumophila* genomic comparisons is cgMLST, which builds on the established SBT method by greatly expanding the number of core loci. To investigate the utility of this method to infer clinical isolate genome source attribution, a matrix of cgMLST allelic distances was generated in the same way that patristic distances were used to build distance-based classifiers. The results from this approach were a slight improvement over the patristic distance-based classifiers, with a higher overall mean F1 score of 0.56, however there were a higher number of false positives, again offering meagre specificity and poor overall classification capacity.

A machine learning classification framework was developed using pan-genome SNP variants to make probabilistic assignments by firstly training models upon variation among environmental isolate genomes to then classify the origins of clinical isolate genomes. To achieve this, an extensive cross-validation framework was established that assessed the performance of various model building parameters (see methods) on the ability for an algorithm to learn upon a portion of environmental isolate genomes and then assign the known classes of the remaining environmental representatives (cross-validation), with the best classification models selected to then learn using the entire training set to make assignments upon the previously unseen clinical isolate genomes.

The application of the machine learning models for the assignment of 113 test set clinical isolate genomes had the greatest classification capacity with 13 out of 20 models achieving an F1-score of 1, indicating perfect sensitivity and specificity. The machine learning method also achieved the greatest overall mean F1 score of 0.70 when evaluating the 14 groups with two or more environmental representatives and 0.74 when applied to all 20 groups. The higher performance of the machine learning modelling approach compared to phylogenomic tree branch length distance and cgMLST allelic distance methods is likely since 1) it considered SNP variation across the pan-genome, 2) it explicitly made use of the underlying sequence composition of the SNP variation and 3) it employed a multivariate approach that modelled the concerted interactions of all input variants. Together, these three aspects of the modelling approach work to make efficient use of the richness of the available SNP allelic variation to achieve greater classification capacity.

False positives were detected with machine learning models ESSEX-A, ESSEX-B and ESSEX-G. Here, the false positives were from other wards in the same hospital, suggesting a sort of ‘cross reactivity’ among nearby locations within a common institution. Despite these false positives, the 74 unrelated clinical isolates were correctly assigned as true negatives by all final models, indicating overall satisfactory negative classification capacity. False negative assignments occurred with models MELB-A, MELB-C, ESSEX-A, ESSEX-B, ESSEX-E and NY-3. In the case of MELB-C and NY-3, previous analyses have identified that there likely exists an issue with the epidemiological source attribution for these outbreak groups, offering a possible explanation for the inability of the models to accurately assign these isolate genomes to their known origins in previous investigations (Buultjens et al., 2017; Raphael et al., 2016).

For MELB-A, the four clinical isolates assigned as false negatives by the machine learning approach were on a branch in the phylogeny that did not contain any environmental isolate genomes from the MELB-A outbreak group, meaning this specific genotype was not represented in the training data. Despite this, all MELB-A clinical isolate genomes were within the top 21% of all clinical isolate genomes when ranked according to decreasing classification probability. This suggests that the modelling approach was able to make use of the level of shared ancestry among all MELB-A isolates to nevertheless provide a useful degree of probability ranking even when that specific genotype was not explicitly represented in the training data. Not dissimilar to what was seen with the MELB-A probability ranking, the classification probabilities for the ESSEX-A, ESSEX-B and ESSEX-G clinical isolate genomes revealed that the known positives for each of these groups were ranked highly despite being less than the standard classification threshold of 0.5. This highlights that alternative probability evaluation frameworks besides classification, such as probability ranking, should be considered for these approaches.

In addition to the use of a ‘one-vs-rest’ classification approach, another notable point of difference with this new method was the use of pan-genome SNPs derived from the reference independent kmer-based method, SKA. Our previous work built models using only variation in core-genome SNPs that were called using read alignment to a reference genome (Buultjens et al., 2017). The consequence of using pan-genome variation was particularly important in this application since the core- genome among the diverse group of 534 *L. pneumophila* isolates is abbreviated, therefore reducing the total amount of SNP diversity. Specifically, the pan-genome alignment provided 258,266 more SNPs than when only core genome variants were considered, equating to addition information to be learnt by multivariate approaches.

The lack of environmental isolates representing a specific MELB-A genotype that was observed exclusively among clinical isolates indicates that the methods used to sample, culture and sequence *L. pneumophila* from environmental sources had failed to adequately capture the true extent of bacterial diversity in that source. Efforts to capture environmental *L. pneumophila* diversity typically involve taking multiple colony picks from environmental samples. While care was taken in this approach to maximise the environmental diversity captured, there evidently was relevant diversity that did not progress to culture isolation and subsequent genome sequencing, presumably due to the limited sensitivity of culture-based methods (Reller, Weinstein, & Murdoch, 2003). Reduced detection of genomic diversity among environmental samples compared to that recovered from clinical specimens has been observed in a previous investigation (Wüthrich et al., 2019). Alternative methods that would likely widen the capture of environmental diversity are shotgun metagenomic or culture independent sequencing approaches that directly sequence all environmental DNA, eliminating the bottleneck of culture (Christiansen et al., 2014; Wéry et al., 2008).

All outbreak groups, apart from MELB-2018, MELB-G and ESSEX-G, had very few numbers of environmental isolate genomes and in some cases just a single genome. Such limited examples of environmental genomic diversity are not optimal and the inclusion of greater numbers of training genomes for each group would likely improve the ability of the models to learn about outbreak specific signatures and make more accurate classifications.

While this study focused on SNP variation, there may be further genomic information among additional variant types such as kmers counted directly from raw reads that may further improve model performance. Such kmer variation has the potential to capture additional genomic variations such as structural variations and copy number differences that were not assessed in this study. Further work may also investigate the specific genomic variants that permit the building of accurate classification models. Such outbreak associated variants may be diagnostic of specific point sources and thus may be informative to understand bacterial genomic responses to certain environmental reservoirs or public health control measures (e.g., different decontamination or biocide practices).

Given the dynamic nature of bacterial populations, routine re-building of the models with newly collected environmental isolates may be required to ensure accuracy as emerging genomic signatures are then learned by the model. Another consideration might be to limit the length of time in which genomes remain in the training database, as older genomic signatures may no longer represent extant *L. pneumophila* in environmental sources as time goes by. Here, a temporal sliding window could be used, as has been implemented in other bacterial genomic investigations (Gorrie et al., 2021).

## CONCLUSION

The advent of highly accessible bacterial genomics has provided a wealth of *L. pneumophila* genomes in publicly assessable databases that are paired with epidemiological information, of which provide the basis to build source attribution classification approaches. Our development of an improved machine learning classification technique now affords models with the ability to call true negatives, offering the previously lacking negative classification capacity. Here we demonstrate that our improved approach provides greater source tracking ability than two widely used methods – phylogenomic trees and cgMLST allelic variation. Given the reported high classification capacity of this improved approach, it is the vision of this work that, soon, future LD public health investigations may make use of such modelling advancements to rapidly pinpoint the correct environmental sources of *L. pneumophila* and reduce the incidence of this preventable disease.

## ACKNOWLEDGEMENTS

We acknowledge the staff of the Health Protection Branch at the Victorian Department of Health for the collection and provision of public health surveillance data used in this study, and their ongoing contribution to the NHMRC Public Health Genomics Partnership.

## REFERENCES

1. Abrams, A. J., & Trees, D. L. (2017). Genomic sequencing of Neisseria gonorrhoeae to respond to the urgent threat of antimicrobial-resistant gonorrhea. Pathogens and disease, 75(4).

2. Bankevich, A., Nurk, S., Antipov, D., Gurevich, A. A., Dvorkin, M., Kulikov, A. S., … Prjibelski, A. D. (2012). SPAdes: a new genome assembly algorithm and its applications to single-cell sequencing. Journal of computational biology, 19(5), 455–477.

3. Bolger, A. M., Lohse, M., & Usadel, B. (2014). Trimmomatic: a flexible trimmer for Illumina sequence data. Bioinformatics, 30(15), 2114–2120.

4. Borchardt, J., Helbig, J., & Lück, P. (2008). Occurrence and distribution of sequence types among Legionella pneumophila strains isolated from patients in Germany: common features and differences to other regions of the world. European Journal of Clinical Microbiology & Infectious Diseases, 27(1), 29–36.

5. Boser, B. E., Guyon, I. M., & Vapnik, V. N. (1992). A training algorithm for optimal margin classifiers. Paper presented at the Proceedings of the fifth annual workshop on Computational learning theory.

6. Breiman, L. (1996). Bagging predictors. Machine learning, 24(2), 123–140.

7. Breiman, L. (2001). Random forests. Machine learning, 45(1), 5–32.

8. Buultjens, A. H., Chua, K. Y., Baines, S. L., Kwong, J., Gao, W., Cutcher, Z., … Tomita, T. (2017). A supervised statistical learning approach for accurate Legionella pneumophila source attribution during outbreaks. Applied and environmental microbiology, 83(21), e01482–01417.

9. Christiansen, M. T., Brown, A. C., Kundu, S., Tutill, H. J., Williams, R., Brown, J. R., … Dave, J. (2014). Whole-genome enrichment and sequencing of Chlamydia trachomatisdirectly from clinical samples. BMC infectious diseases, 14(1), 1–11.

10. David, S., Afshar, B., Mentasti, M., Ginevra, C., Podglajen, I., Harris, S. R., … Parkhill, J. (2017). Seeding and establishment of Legionella pneumophila in hospitals: implications for genomic investigations of nosocomial Legionnaires’ disease. Clinical Infectious Diseases, 64(9), 1251–1259.

11. David, S., Rusniok, C., Mentasti, M., Gomez-Valero, L., Harris, S. R., Lechat, P., … Ma, L. (2016). Multiple major disease-associated clones of Legionella pneumophila have emerged recently and independently. Genome research, 26(11), 1555–1564.

12. Didelot, X., & Wilson, D. J. (2015). ClonalFrameML: efficient inference of recombination in whole bacterial genomes. PLoS computational biology, 11(2), e1004041.

13. Fields, B. S., Benson, R. F., & Besser, R. E. (2002). Legionella and Legionnaires’ disease: 25 years of investigation. Clin Microbiol Rev, 15(3), 506–526.

14. Géron, A. (2019). Hands-on machine learning with Scikit-Learn, Keras, and TensorFlow: Concepts, tools, and techniques to build intelligent systems: “O’Reilly Media, Inc.”.

15. Goldberg, B., Sichtig, H., Geyer, C., Ledeboer, N., & Weinstock, G. M. (2015). Making the leap from research laboratory to clinic: challenges and opportunities for next-generation sequencing in infectious disease diagnostics. MBio, 6(6), e01888–01815.

16. Gorrie, C. L., Da Silva, A. G., Ingle, D. J., Higgs, C., Seemann, T., Stinear, T. P., … Sherry, N. L. (2021). Key parameters for genomics-based real-time detection and tracking of multidrug-resistant bacteria: a systematic analysis. The Lancet Microbe, 2(11), e575–e583.

17. Gorzynski, J., Wee, B., Llano, M., Alves, J., Cameron, R., McMenamin, J., … Fitzgerald, J. R. (2022). Epidemiological analysis of Legionnaires’ disease in Scotland: a genomic study. The Lancet Microbe, 3(11), e835–e845.

18. Graham, R., Doyle, C., & Jennison, A. (2014). Real-time investigation of a Legionella pneumophila outbreak using whole genome sequencing. Epidemiology & Infection, 142(11), 2347–2351.

19. Harris, S. R. (2018). SKA: Split kmer analysis toolkit for bacterial genomic epidemiology. bioRxiv, 453142.

20. Harrison, T., Afshar, B., Doshi, N., Fry, N., & Lee, J. (2009). Distribution of Legionella pneumophila serogroups, monoclonal antibody subgroups and DNA sequence types in recent clinical and environmental isolates from England and Wales (2000–2008). European Journal of Clinical Microbiology & Infectious Diseases, 28(7), 781–791.

21. Ingle, D. J., Howden, B. P., & Duchene, S. (2021). Development of phylodynamic methods for bacterial pathogens. Trends in Microbiology, 29(9), 788–797.

22. Krøvel, A. V., Bernhoff, E., Austerheim, E., Soma, M. A., Romstad, M. R., & Löhr, I. H. (2022). Legionella pneumophila in Municipal Shower Systems in Stavanger, Norway; A Longitudinal Surveillance Study Using Whole Genome Sequencing in Risk Management. Microorganisms, 10(3), 536.

23. Kwong, J. C., Mercoulia, K., Tomita, T., Easton, M., Li, H. Y., Bulach, D. M., … Howden, B. P. (2016). Prospective whole-genome sequencing enhances national surveillance of Listeria monocytogenes. Journal of clinical microbiology, 54(2), 333–342.

24. Lück, C., Fry, N. K., Helbig, J. H., Jarraud, S., & Harrison, T. G. (2013). Typing methods for Legionella. In Legionella (pp. 119–148): Springer.

25. McAdam, P. R., Vander Broek, C. W., Lindsay, D. S., Ward, M. J., Hanson, M. F., Gillies, M., … Fitzgerald, J. R. (2014). Gene flow in environmental Legionella pneumophila leads to genetic and pathogenic heterogeneity within a Legionnaires’ disease outbreak. Genome biology, 15(11), 1–10.

26. Mercante, J. W., & Winchell, J. M. (2015). Current and emerging Legionella diagnostics for laboratory and outbreak investigations. Clinical microbiology reviews, 28(1), 95–133.

27. Moran-Gilad, J., Prior, K., Yakunin, E., Harrison, T., Underwood, A., Lazarovitch, T., … Agmon, V. (2015). Design and application of a core genome multilocus sequence typing scheme for investigation of Legionnaires’ disease incidents. Eurosurveillance, 20(28), 21186.

28. Paradis, E., Claude, J., & Strimmer, K. (2004). APE: analyses of phylogenetics and evolution in R language. Bioinformatics, 20(2), 289–290.

29. Petzold, M., Prior, K., Moran-Gilad, J., Harmsen, D., & Lück, C. (2017). Epidemiological information is key when interpreting whole genome sequence data–lessons learned from a large Legionella pneumophila outbreak in Warstein, Germany, 2013. Eurosurveillance, 22(45), 17–00137.

30. Price, M. N., Dehal, P. S., & Arkin, A. P. (2009). FastTree: computing large minimum evolution trees with profiles instead of a distance matrix. Molecular biology and evolution, 26(7), 1641–1650.

31. Qin, T., Zhang, W., Liu, W., Zhou, H., Ren, H., Shao, Z., … Xu, J. (2016). Population structure and minimum core genome typing of Legionella pneumophila. Scientific reports, 6(1), 1–10.

32. Raphael, B. H., Baker, D. J., Nazarian, E., Lapierre, P., Bopp, D., Kozak-Muiznieks, N. A., … Musser, K. A. (2016). Genomic resolution of outbreak-associated Legionella pneumophila serogroup 1 isolates from New York State. Applied and environmental microbiology, 82(12), 3582–3590.

33. Reller, L. B., Weinstein, M. P., & Murdoch, D. R. (2003). Diagnosis of Legionella infection. Clinical Infectious Diseases, 36(1), 64–69.

34. Reuter, S., Harrison, T. G., Köser, C. U., Ellington, M. J., Smith, G. P., Parkhill, J., … Török, M. E. (2013). A pilot study of rapid whole-genome sequencing for the investigation of a Legionella outbreak. BMJ open, 3(1), e002175.

35. Ricci, M. L., Fillo, S., Ciammaruconi, A., Lista, F., Ginevra, C., Jarraud, S., … Lindsay, D. (2022). Genome analysis of Legionella pneumophila ST23 from various countries reveals highly similar strains. Life science alliance, 5(6).

36. Rousseau, C., Ginevra, C., Simac, L., Fiard, N., Vilhes, K., Ranc, A.-G., … Campese, C. (2022). A Community Outbreak of Legionnaires’ Disease with Two Strains of L. pneumophila Serogroup 1 Linked to an Aquatic Therapy Centre. International Journal of Environmental Research and Public Health, 19(3), 1119.

37. Sánchez-Busó, L., Guiral, S., Crespi, S., Moya, V., Camaró, M. L., Olmos, M. P., … Vanaclocha, H. (2016). Genomic investigation of a legionellosis outbreak in a persistently colonized hotel. Frontiers in microbiology, 6, 1556.

38. Schoonmaker-Bopp, D., Nazarian, E., Dziewulski, D., Clement, E., Baker, D. J., Dickinson, M. C., … Lapierre, P. (2021). Improvements to the Success of Outbreak Investigations of Legionnaires’ Disease: 40 Years of Testing and Investigation in New York State. Applied and environmental microbiology, 87(16), e00580–00521.

39. Schwake, D. O., Garner, E., Strom, O. R., Pruden, A., & Edwards, M. A. (2016). Legionella DNA markers in tap water coincident with a spike in Legionnaires’ disease in Flint, MI. Environmental Science & Technology Letters, 3(9), 311–315.

40. Sintchenko, V., & Holmes, E. C. (2015). The role of pathogen genomics in assessing disease transmission. Bmj, 350.

41. Wéry, N., Bru-Adan, V., Minervini, C., Delgénes, J.-P., Garrelly, L., & Godon, J.-J. (2008). Dynamics of Legionella spp. and bacterial populations during the proliferation of L. pneumophila in a cooling tower facility. Applied and environmental microbiology, 74(10), 3030–3037.

42. Wüthrich, D., Gautsch, S., Spieler-Denz, R., Dubuis, O., Gaia, V., Moran-Gilad, J., … Tschudin-Sutter, S. (2019). Air-conditioner cooling towers as complex reservoirs and continuous source of Legionella pneumophila infection evidenced by a genomic analysis study in 2017, Switzerland. Eurosurveillance, 24(4), 1800192.

43. Yu, V. L., Plouffe, J. F., Pastoris, M. C., Stout, J. E., Schousboe, M., Widmer, A., … Paterson, D. L. (2002). Distribution of Legionella species and serogroups isolated by culture in patients with sporadic community-acquired legionellosis: an international collaborative survey. The Journal of infectious diseases, 186(1), 127–128.

